# A Comparison of the Costs and Benefits of Bacterial Gene Expression

**DOI:** 10.1101/038851

**Authors:** Morgan N. Price, Kelly M. Wetmore, Adam M. Deutschbauer, Adam P. Arkin

## Abstract

To study how a bacterium allocates its resources, we compared the costs and benefits of most of the proteins in *Escherichia coli* K-12 during growth in minimal glucose medium. Proteins that are important for fitness are usually highly expressed, and 95% of these proteins are expressed at above 13 parts per million (ppm). Conversely, proteins that do not measurably benefit the host tend to be weakly expressed, with a median expression of 13 ppm. In aggregate, genes with no detectable benefit account for 31% of protein production, or about 22% if we correct for genetic redundancy. Although some of the apparently unnecessary expression could have subtle benefits in minimal glucose medium, the majority of the burden is due to genes that are important in other conditions. We propose that over 10% of the cell’s protein is “on standby” in case conditions change.

## Introduction

The typical bacterial genome encodes thousands of proteins, and many of these proteins are not beneficial for growth at any given time. For example, the model bacterium *Escherichia coli* K-12 preferentially utilizes glucose. Its genome encodes hundreds of genes that enable it to utilize other carbon sources, but these genes will not be beneficial if glucose is available. Furthermore, the activity of many proteins can be detrimental, as the loss of many genes confers a measurable growth advantage in some conditions (1; 2; 3; 4; 5).

Expressing an unnecessary protein should reduce the growth rate even if the protein’s activity is harmless. In theoretical models of microbial growth, useless protein causes a reduction in fitness (or the relative growth rate) equal to the fraction of all protein that is useless (6; 7) or a small multiple of this (8). In laboratory environments, the measured fitness cost of a useless and harmless protein is about 1-2 times the fraction of protein (9; 8; 10).

Bacterial proteins are typically expressed at 3-21 parts per million of the protein mass of a cell (data of (11), 25th-75th percentile). Although a cost of 3 ppm may seem small, it should be significant for evolution. The effective population sizes (N_e_) for bacteria are estimated at around 10^6^ or 10^7^ (12), and under the nearly-neutral theory of molecular evolution, this implies that alleles that increase fitness by just 1 ppm should predominate over evolutionary time (N_e_ · *s* > 1, where *s* is the selection coefficient; see (13)).

Given the high cost of unnecessary expression, bacteria should evolve to allocate their expression of protein to genes that are important for growth or survival. Several recent studies examined the concentrations of proteins in bacteria as a resource allocation problem. In *E. coli*, proteins that are regulated by the growth rate account for about half of the protein mass (14), and the total expression of many functional categories of proteins is correlated with the growth rate (15). However, the importance of these proteins for growth or fitness was not examined. In *Bacillus subtilis*, the concentrations of most enzymes can be explained by their theoretically-optimal flux (16); this study included metabolic enzymes, ribosomes, and chaperones, but did not include most of the genes in the genome. Also, all of these studies relied on peptide mass-spectrometry (“proteomics”), and so they focused on relatively highly-expressed proteins. These studies reported abundances for just 18-55% of the proteins in the genome.

Here, we compare the costs and benefits of 86% of the proteins in *E. coli* K-12 during growth in a minimal glucose medium. To measure protein production or cost, we used ribosomal profiling data (11), which allows us to study weakly-expressed proteins. To measure the benefit of each protein, or its importance for growth, we used a barcoded library of about 150,000 transposon mutants (17) as well as information from individual knockout strains (18; 19). We found that 96% of protein-coding genes that had mutant phenotypes were expressed at above 10 ppm of protein mass or above 40 monomers per cell, and their median expression was 205 ppm. In contrast, genes that did not have a measurable impact on fitness had a median expression of 13 ppm. Overall, genes that were not important for fitness accounted for 31% of protein production by mass, but some of these proteins are isozymes or are otherwise expected to be redundant. Once we correct for genetic redundancy, we estimate that in this condition, 22% of protein production is unnecessary. Many of these proteins are only expected to be important in other conditions, given their known functions. Indeed, by examining a large compendium of genome-wide fitness assays, we found that the majority of this unnecessary expression is for proteins that have significant phenotypes in other conditions. We propose that most of the 22% burden is for proteins that are “on standby” in case conditions change.

## Results

### Comparison of ribosomal profiling to mutant phenotypes

To compare the costs and benefits of gene expression, we studied *E. coli* K-12 growing at 37°C in MOPS minimal glucose media. We obtained ribosomal profiling data from Li and colleagues (11) and we use the fraction of protein expression (weighted by the length of the protein) to estimate the cost of expression. The ribosomal profiling data should be accurate to within 2-fold for most genes, as the data from two halves of a gene, or for two proteins in an equimolar complex, tend to be consistent within this range (11). Also, Li and colleagues report that their quantitation is accurate for genes with over 128 reads, which corresponds to roughly 1 ppm of expression. Genes with fewer reads are probably expressed at under 1 ppm. For a protein of average size, 1 ppm corresponds to about 6 monomers being produced per cell cycle, as in these conditions there are 5.6 million protein monomers per cell (11).

To measure the importance of each protein for growth, we used a pooled library of about 150,000 transposon mutants with DNA barcodes (17). We grew the pooled mutants in MOPS minimal glucose media for 12 generations and we assayed the abundance of each mutant before and after growth by amplifying the DNA barcodes, sequencing them, and counting them (17). Because mutants that have a strong growth defect in rich media will be missing from our library, we also used a list of 286 essential (or nearly-essential) proteins from PEC (19) and growth data on individual deletion strains from the Keio collection (18). From PEC, we identified 286 essential proteins; we classified non-essential genes whose mutants had a significant change in abundance during our pooled assay as either important for fitness (294 genes) or detrimental to fitness (25 genes); we classified 25 other genes with low coverage in our mutant pools as important for fitness because mutants grew poorly in both minimal MOPS media and LB (18); we classified proteins with non-significant changes in mutant abundance of above 3% per generation (48 genes) or with insufficient coverage (449 genes) as ambiguous; and we classified the remaining 2,944 proteins as having no phenotype. Based on a negative control in which we examined the differences between two independently-grown samples of the pool of mutants, we expect around 1 false positive among the genes with significant phenotypes (see Methods). The genes with no phenotype probably affect growth by less than 5% per generation, because of the genes with estimated effects of 4%-5% per generation, 28 of 39 (72%) were statistically significant. The phenotypic classification of the genes is included in Supplementary Table S1.

As shown in Figure 1A, genes that have a phenotype are much more likely to be highly expressed, but even genes with no phenotype are usually expressed at significant levels, with a median expression of 13 ppm. Also note that 13 ppm is far above the level at which the ribosomal profiling data becomes noisy, which is about 1 ppm. 81% of proteins with no phenotype are expressed at above 1 ppm.

### Most genes that affect fitness are well expressed

Proteins with detectable benefits in minimal glucose medium tended to be well expressed in this condition, with 96% of these genes being expressed at 10 ppm or more. Similarly, 23 of 25 of proteins that were detrimental to fitness (92%) were expressed at 10 ppm or more, which makes sense because they should be expressed at a significant level in order to have a measurable negative impact on the cell. (For gene sizes in the 25th-75th percentile, 10 ppm corresponds to 30-70 monomers per cell.)

All of the essential proteins (19) were expressed at above 5 ppm, except for the putative protein YceQ, which had no ribosomal profiling reads at all. YceQ also lacked ribosomal profiling reads in two other growth conditions (11); YceQ lacks homology to any other protein; the open reading frame is disrupted in some strains of *E. coli*; and we identified transposon insertions within *yceQ* ((17); Supplementary Figure S1). It appears that *yceQ* does not encode a protein. Deletion of the entire *yceQ* region may not be possible because it contains the promoter of the essential gene *rne* (Supplementary Figure S1). The other weakly-expressed essential proteins (below 10 ppm) were PrmC and MreD, at around 30 and 60 monomers per cell, respectively. (PrmC, formerly known as HemK, was listed as essential by (19); it is not entirely essential but a mutant has severely reduced growth (20).)

**Figure 1.**
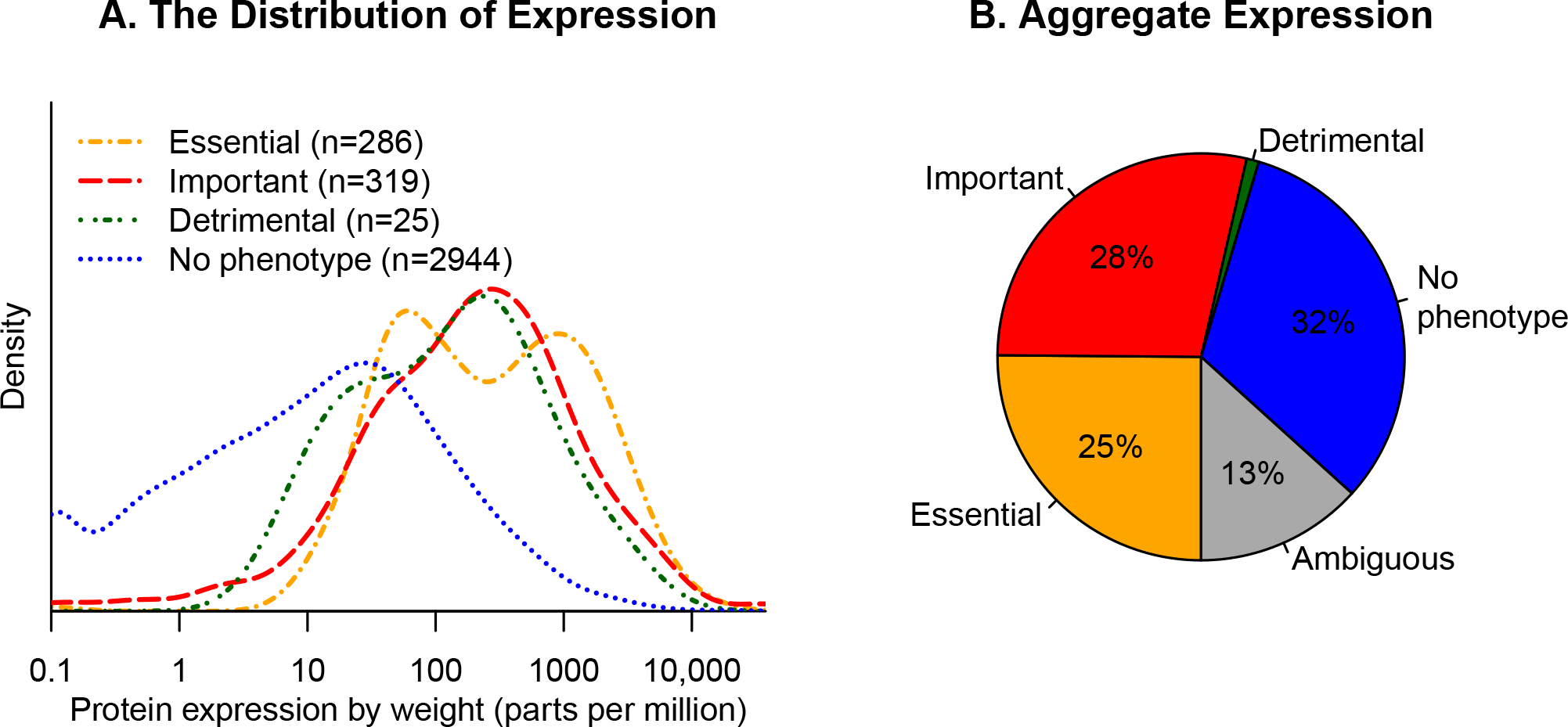
**The production of proteins versus their importance for growth.** (A) For each class of gene, we show the distribution of protein expression, in parts per million of amino acids (*x* axis, log scale). Proteins with little or no expression are shown at 0.1 ppm. (B) The aggregate expression of each class of gene.

Some very weakly-expressed non-essential genes were identified as being important for fitness, including 6 proteins with expression of under 1 ppm of monomers or roughly 6 copies per cell. These proteins were ArpA, WcaE, YahL, YbfK, YdbD, and YnbB. WcaE is believed to be a glycosyltransferase that is involved in the biosynthesis of colanic acid, an exopolysaccharide; little is known about the function of the other proteins (21). We are not sure how these proteins could have a measurable effect at 6 copies per cell unless they are regulatory proteins. Another 7 genes were important for fitness despite weak expression of 1-2 ppm of monomers (6-12 copies per cell), including three proteins that are involved in the uptake of iron via the siderophore enterobactin (FepD, FepG, and Fes). Differences between the genetic backgrounds of the two data sets, or subtle differences in growth conditions, might lead to these rare discrepancies (see Methods). We also wondered if these proteins might be important for the transition to growth in this condition, rather than during exponential growth (which is when protein production was measured), but this does not seem to be the case (see Methods).

Overall, we find that almost all proteins are expressed at above 10 ppm of monomers (or roughly 50 monomers per cell) when they have a significant effect on growth. This threshold accounts for 95% of the genes with phenotypes (599 of 630).

### High expression of many proteins with no expected benefit

31% of total expression (by mass) was due to the 2,944 proteins with no measurable impact on fitness (Figure 1B). As shown in Figure 1A, the distribution of expression is quite skewed, so most of this 31% is due to a few hundred well-expressed genes. We decided to focus on 287 proteins were expressed at above 200 ppm and had no measurable impact on growth; these account for 24% of total protein production. Proteomics data (15) confirms that most of these “unnecessary” proteins are highly expressed (see Methods).

**Table I.** **The 20 most highly-expressed genes, by fraction of amino acids, that have no measurable impact on growth.**

| Gene | Fraction | Group | Description |
| --- | --- | --- | --- |
| *ompF* | 2.0% | Key/redundant | outer membrane porin 1a |
| *ompC* | 1.5% | Key/redundant | outer membrane porin protein C |
| *livJ* | 0.6% | Nutrient | leucine/isoleucine/valine transporter subunit |
| *ompT* | 0.6% | Stress | outer membrane protease VII |
| *ahpC* | 0.5% | Key/redundant | alkyl hydroperoxide reductase, C22 subunit |
| *aceA* | 0.5% | Central | isocitrate lyase (glyoxylate shunt) |
| *pflB* | 0.4% | Anaerobic | pyruvate formate lyase I |
| *aceB* | 0.3% | Central | malate synthase (glyoxylate shunt) |
| *zinT* | 0.3% | Stress | cadmium-induced cadmium binding protein |
| *sodA* | 0.3% | Key/redundant | superoxide dismutase, manganese |
| *adhE* | 0.3% | Anaerobic | aldehyde-alcohol dehydrogenase |
| *pyrI* | 0.3% | – | aspartate carbamoyltransferase regulatory subunit |
| *ompX* | 0.3% | – | outer membrane protein X |
| *oppA* | 0.3% | Nutrient | oligopeptide transporter subunit |
| *pntA* | 0.2% | Central | NAD(P) transhydrogenase subunit alpha |
| *sodB* | 0.2% | Key/redundant | superoxide dismutase, Fe |
| *pykF* | 0.2% | Key/redundant | pyruvate kinase |
| *metQ* | 0.2% | Nutrient | DL-methionine transporter subunit |
| *dppA* | 0.2% | Nutrient | dipeptide transporter |
| *tpx* | 0.2% | Stress | thiol peroxidase |

We show the 20 most highly-expressed proteins with no mutant phenotype in Table 1. Six of these genes are involved in key processes that are important for growth but the knockout lacks a phenotype because of genetic redundancy. For example, the top two proteins, OmpF and OmpC, are the two major outer membrane porins. Presumably, some expression of porins is necessary for the movement of nutrients through the outer membrane, but deleting either of these individually has little effect. Similarly, AhpC reduces hydrogen peroxide, which is a toxic byproduct of the aerobic electron transport chain, but so do KatE and KatG. If all three genes are disabled, then under aerobic conditions, toxic hydrogen peroxide will accumulate and growth will be inhibited (22). SodA and SodB are redundant isozymes of superoxide dismutase and eliminate another toxic byproduct of oxygen utilization. A strain lacking both SodA and SodB cannot grow aerobically in a minimal glucose medium (23). Finally, PykF is one of two isozymes of pyruvate kinase, which (in reverse) is an ATP-forming step in glycolysis. Again, these isozymes are likely redundant.

Of the remaining 14 highly-expressed proteins with no mutant phenotype, 12 are only expected to be important for growth in other conditions. High expression of these proteins might nevertheless be selected for, just in case growth conditions change (4). For example, AceA and AceB are involved in central metabolism – they encode the glyoxylate shunt – but a metabolic model predicts that they are not required for optimal growth on glucose (24), and ^13^C labeling studies suggest that they do not carry flux in this condition (25; 26). PntA (the pyridine nucleotide dehydrogenase *α* subunit) is also involved in central metabolism and is predicted to be dispensible for optimal growth on glucose (24), although it is not clear whether this prediction is correct. (A strain that lacked both subunits *(*pntAB*^−^)* was reported to have a 30% reduction in growth rate on glucose (25), while we observed a defect of just 2% in mutants of either *pntA* or *pntB*, and this defect was not statistically significant. The discrepancy could be due to differences in growth conditions.) Similarly, in *B. subtilis*, some enzymes in central metabolism are highly expressed even when they carry no flux (16). LivJ, OppA, MetQ, and DppA are involved in the uptake of amino acids or short peptides, which are not present in our media. PflB and AdhE are probably important for growth in anaerobic conditions. OmpT, ZinT, and Tpx are involved in responses to stresses that were not present in our experiment. The two remaining proteins are a regulatory subunit of an important enzyme (PyrI) and and an outer membrane protein whose function is not well understood (OmpX).

To more systematically examine the highly-expressed and “unnecessary” proteins, we examined the EcoCyc entries for all 287 highly-expressed genes that lack phenotypes (Supplementary Table S2). 106 of these 287 proteins are expected to be important in other conditions, as they are involved in utilizing alternate nutrients (66 proteins), stress resistance (24 proteins), parts of central metabolism that are dispensible in our condition (11 proteins), or anaerobic growth (5 proteins). These 106 unnecessary proteins account for 11.4% of total protein production. Another 60 of the 287 proteins are involved in key processes that are expected to be important for growth in our condition, but are redundant because of isozymes (as with SodA and SodB above) or more indirect redundancy (see appendix). These 60 genetically-redundant proteins account for 6.8% of protein production. Another 10 proteins are involved in key processes and have little phenotype, even though they are not redundant as far as we know: these include 4 non-essential components of essential complexes and 2 proteins involved in the modification of tRNA or rRNA (see appendix). These proteins might have fine-tuning or regulatory roles. Just 8 of the 287 proteins are characterized transcriptional regulators (27). The majority of the remaining genes are poorly characterized: 62/103 (60%) have names that begin with *y*.

If we assume that the proportion of “unnecessary” expression that is due to genetic redundancy is similar for moderately-expressed proteins as it is for highly-expressed proteins (6.8/24.1 = 28%), then we estimate that 31% · (1 − 0.28) = 22% of protein production is unnecessary in this growth condition. More conservatively, if we consider only the proteins of known function that are not expected to be important, then we estimate that 11.4/24.1 · 31% = 15% of protein production is unnecessary.

### Highly-expressed genes that are not important for fitness often have phenotypes in other conditions

If much of unnecessary protein production is due to preparation for other conditions, then the highly-expressed yet unnecessary proteins should be important for fitness in other conditions. So, we asked if these genes have phenotypes in a compendium of 162 fitness experiments for *E. coli* ((17); M. N. Price *et al.*, in preparation; http://fit.genomics.lbl.gov/). This compendium includes growth in 29 different carbon sources, growth in 16 different nitrogen sources, growth in the presence of 35 different antibiotics or biocides, and motility on an agar plate. The compendium covers 2,944 proteins that do not have a phenotype in minimal glucose media, and 722 of these are important for growth in other conditions or for motility. As shown in Figure 2, proteins that are not important in minimal glucose media are much more likely to be highly expressed if they have phenotypes in other conditions, with a median expression of 45 ppm instead of 7 ppm *(P < *10^−15^, Wilcoxon rank sum test). In total, the 722 proteins that are not important for fitness but have phenotypes in other conditions account for more of protein production in minimal glucose media (18%) than the 2,222 proteins that do not have any phenotypes at all (14%).

A caveat is that some of the genes with measurable phenotypes in artifical conditions might be more subtly useful in nature. For example, some form of cellular damage might occur at low rates under natural conditions, so that the repair genes have subtle benefits. Yet during growth in the presence of an inhibitor that creates this type of damage, these fine-tuning genes could have large benefits. We thought that this caveat would be less likely to be relevant for carbon sources or nitrogen sources: *E. coli* would probably not be able to consume these nutrients unless they were sometimes important in nature. So we considered each carbon or nitrogen source experiment individually, and asked whether proteins that were important for utilizing that nutrient (but not glucose) where more highly expressed than the typical protein (in minimal glucose media). Also, we considered only experiments with 5 or more important proteins. In 82 of 96 experiments, the median protein that was important for fitness (in the condition but not in glucose) was expressed at least 3-fold higher than the median protein (in glucose). Thus, we propose that much of the “unnecessary” expression represents preparation for changing conditions.

**Figure 2.**
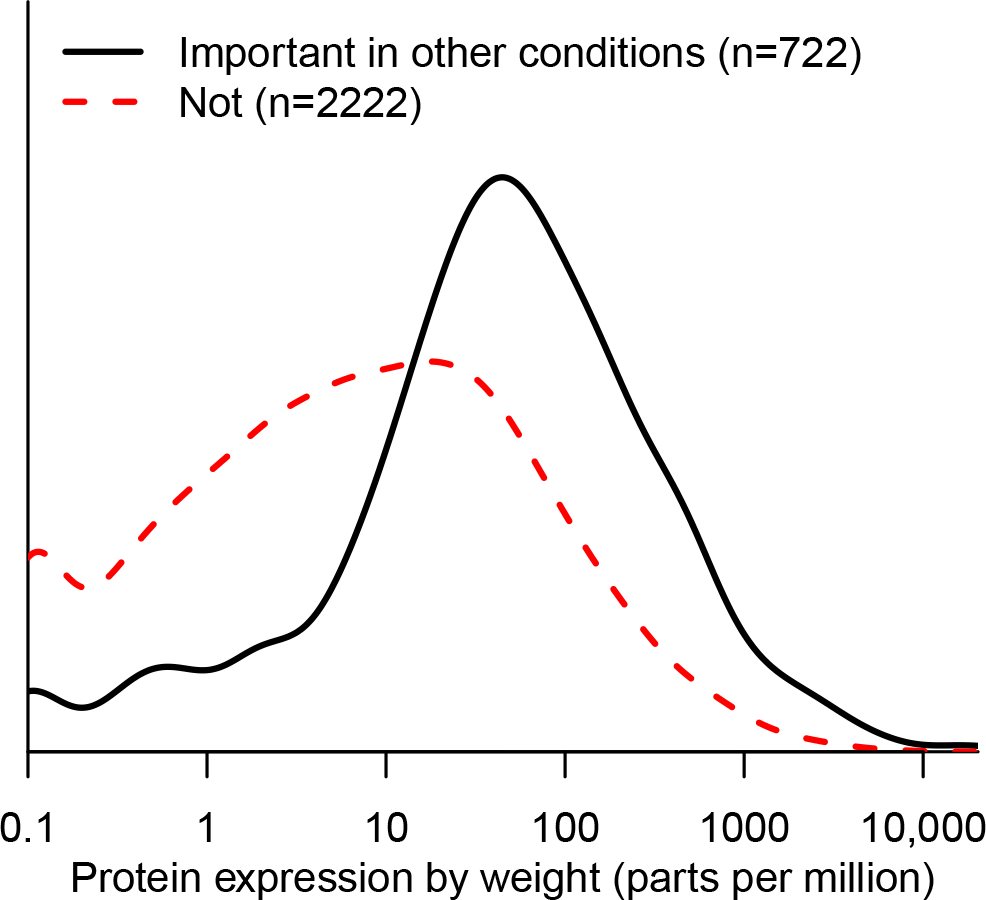
**The production of “unnecessary” proteins versus their importance in other conditions.** Only genes that were not important for fitness in minimal glucose media are included. The *x* axis is as in Figure 1A.

## Discussion

### Do “unnecessary” proteins have subtle benefits?

A major caveat in our results is that the “unnecessary” proteins might have benefits that are too subtle for us to measure. Our assay is sensitive to a loss of fitness due to disabling a gene of about 4% per generation, which would give a log_2_ ratio of −0.7 after 12 generations. But natural selection could maintain the expression of proteins with far smaller benefits.

We relied on our understanding of *E. coli’s* physiology to argue that many of these proteins are unlikely to provide a subtle benefit. Highly-expressed proteins that are unlikely to be beneficial were involved in steps in central metabolism that do not carry flux (such as *aceAB*), in anaerobic growth, or in the utilization of nutrients that were not provided in the growth medium.

A few of the “unnecessary” proteins might be involved in the salvage of components that leak out of the cell. For example, of the 66 highly-expressed and “unnecessary” proteins for utilizing alternate nutrients, 16 are involved in the uptake of amino acid or peptides, and these account for 2% of total protein. Although neither amino acids nor peptides were present in our media, they might be released by the cells. For example, peptides might be released after the degradation of misfolded periplasmic proteins by DegP (also known as HtrA) or OmpT, as both of these proteases are highly expressed in our growth condition. But we doubt that the secretion of amino acids or peptides is sufficient to justify an investment of 2% of protein in recovering them. A similar scenario arises with glutathione, which can be secreted to a concentration of over 100 *μ*M by *E. coli* while growing in minimal glucose media (28). (The secretion of glutathione may be a consequence of the domestication of *E. coli* K-12, as it was not observed with clinical isolates (29).) The GsiB protein (formerly known as YliB) accounts for 0.02% of protein and is involved in the reuptake of the secreted glutathione (28). Although we did not observe a benefit for *gsiB* — we estimated a +1% per generation change in mutant abundance per generation, which was not statistically significant — it is possible that it has a tiny benefit that is too small for us to observe.

Overall, we cannot be certain of the lack of benefit for any one of these genes, but it does not seem plausible that the expression of most of these genes is beneficial.

### Adaptive just-in-case expression of many genes

Instead, we argue that the expression of many of these proteins is due to preparation for other conditions. This was supported by the observation that many of the “unnecessary” and highly-expressed proteins are measurably important for fitness in other conditions.

The high expression of genes that are important in other conditions need not imply that those genes are constitutively expressed. Of the 227 highly-expressed genes with no measurable phenotype in glucose minimal media and no expectation of genetic redundancy, 45% are known to be regulated by one or more transcription factors (27). This is similar to the rate for all genes (38%). Because many of the transcription factors in *E. coli* K-12 are still poorly characterized, the true proportion could be much higher.

High unnecessary expression of regulated genes may seem paradoxical, but intuitively, if the good times are not likely to last, then it is adaptive to express these genes at significant (but not maximal) levels: turning them on only when needed could lead to a long lag in growth (30; 31; 32). Alternatively, the high expression of these genes in artificial conditions could reflect their regulation by signals that are not directly related to their function (4), and such high levels of unnecessary expression might not occur in natural conditions.

### The cost of being a generalist

We estimate that 22% of *E. coli’s* protein production in minimal glucose medium is due to genes that are not important for fitness in this condition (after correcting for genetic redundancy). And we observed that over half (57%) of this expression was due to genes that are important for fitness in other conditions. This implies that, in minimal glucose media, *E. coli* invests 22% · 57% ≈ 13% of protein to prepare for other conditions. Because the fitness cost of useless protein is at least as large as the fraction of protein (6; 8; 9; 10), this burden could reduce *E. coli’s* growth rate by 13%.

To test if this kind of burden occurs in other microbes, we compared ribosomal profiling data (33) and homozygous mutant data (34) for budding yeast *Saccharomyces cerevisiae* growing in rich media. We found that about 25% of protein production is for genes that are not important for fitness. We suspect that most microbes invest significantly in the expression of proteins that are “on standby” in case conditions change.

The high aggregate cost of unnecessary expression suggests that it might be possible to engineer strains with reduced genomes that will grow faster or more efficiently. For example, Posfai and colleagues constructed a strain of *E. coli* K-12 with 42 deletions that removed 14% of the genome (35). We estimate that deleting these genes saved 2.6% of protein production. (We ignored any regulatory effects of the deletions.) But, Posfai and colleagues also removed six of the genes that we identified as being important for fitness (*yagM, ydbD, ynbB, wcaE, yfdI/gtrS,* and *yfjI*), which may explain why they did not observe any improvement in the growth rate.

Another aspect of being a generalist is the need for gene regulation. Among the 186 homomeric transcription factors that have been characterized in *E. coli* (27). the typical expression was 10 ppm to 60 ppm (25th to 75th percentile), and their total expression is 1.8%. Thus, the total cost of gene regulation does not seem to be that high. However, the cost of an individual regulator would be significant during evolution, which might cause rarely-needed capabilities to be lost from most members of a population (36).

### Proteins with tiny benefits will not be maintained

We propose a minimum threshold for the benefit of a protein, on the assumption that it will not be maintained by natural selection unless the benefit exceeds the cost. We found that 96% of proteins with a detectable fitness advantage had a cost of above 10 ppm. Similarly, in *S. cerevisiae* growing in rich media, 89% of proteins with a detectable fitness advantage are expressed at 10 ppm or higher (combining (34; 33)). It is possible that a protein with subtle benefits might not require such high expression, but we found that the median protein without a measurable phenotype was still expressed at 13 ppm. So, we propose that proteins with benefits of 10 ppm or less will be selected against. Although a benefit of 10 ppm might seem small, such tiny benefits are sometimes considered in evolutionary theory, such as in a recent model of selection for genetic redundancy (37). We argue that evolutionary models that rely on such subtle benefits of a gene are not realistic because of the cost of protein production.

### The challenge of detecting the cost of unnecessary expression

Although an unnaturally highly-expressed gene can cause a measurable reduction in the growth rate (8; 9; 10), it is challenging to measure the reduction in the growth rate due to unnecessary expression at natural levels. In minimal glucose media, many of the highly and “unnecessarily” expressed proteins account for less than 0.1% of protein each, which implies that it would take over 100 generations to see a 10% increase in the relative abundance of a deletion strain. An experiment of this length is risky because of secondary mutations. The most highly-expressed “unnecessary” protein was just 0.6% of protein (LivJ, see (Table 1).

Would the cost of expressing unnecessary protein should be detectable in laboratory evolution experiments? A cost of 0.6% implies that it would take 115 generations for the abundance of a beneficial mutant to increase by even two-fold. If the mutant is initially very rare, then it still will not be detectable. Furthermore, in laboratory evolution experiments with bacteria, strongly beneficial mutations are common, which leads to “hitchhiking” — if a strongly-beneficial mutation happens to arise in a genetic background that contains mildly-deleterious variants, then those deleterious variants will increase in abundance. Thus, hitchhiking weakens the impact of selection on variants with more subtle effects. In theory, variants become effectively neutral if they are under selection that is an order of magnitude weaker than that of the strongly-beneficial mutations (38). Because beneficial mutations often have an advantage of several percent per generation or more (39), the loss of an unnecessary protein that is expressed at 0.3% would be effectively neutral.

In practice, it appears that some unnecessary catabolic capabilities are lost during the evolution of *E. coli* in the laboratory in a glucose medium, but most remain, even after 50,000 generations (40). Losses occur at a higher rate in strains with elevated mutation rates, which suggests that many of the losses are effectively neutral (40). Also, because many of the losses of catabolic capabilities in the mutator strains are rescued by a reduction in temperature, these losses are probably due to mutations that cause proteins to misfold, rather than mutations that reduce unnecessary expression (40). A few catabolic capabilities were lost in multiple lines, which seems to reflect selection against the deleterious activity of a protein, rather than selection on the cost of making an unnecessary protein. For example, during the growth of *E. coli* on glucose, there is strong selection (1-2% per generation) for the loss of D-ribose utilization (41). This seems far too strong to be explained by the cost of expressing the ribose utilization operon, which we estimate at 0.03%. Instead, it appears that the activity of these proteins is detrimental. More broadly, many genes are detrimental to fitness in some conditions (1; 2; 3; 4; 5), and these are the genes that we expect will be lost or down-regulated during laboratory evolution experiments, regardless of their expression levels.

## Conclusions

Proteins that are important for fitness are highly expressed, but many of the highly-expressed proteins are not important for fitness. Some of this is due to genetic redundancy, but most of these proteins are important for fitness in other conditions, so we propose that they are expressed because of the possibility of a rapid change in conditions. In aggregate, this preparation accounts for around 13% of the protein in *E. coli* growing in minimal glucose media. The bulk of this investment is due to around 200 highly-expressed proteins, but most other proteins that are not important for fitness are still expressed at detectable levels and have significant costs during evolution.

## Materials and Methods

### Measuring mutant fitness in minimal glucose medium

We used a collection of 152,018 randomly-barcoded transposon mutants that were derived from *E. coli* strain BW25113 (17). The pool of mutants was recovered from the freezer by growing it in rich medium (LB) until OD_600_ = 1 and pelleted, and an initial sample was collected. The remaining cells were washed and inoculated into two different 2 liter flasks, each with 200 mL of MOPS minimal medium (Teknova) supplemented with 2 g/L D-glucose, at an initial OD = 0.02. (MOPS includes inorganic salts as well as 3-(N-morpholino)propanesulfonic acid and tricine as buffering agents, but does not include any vitamins.) The cells grew aerobically at 37°C until late exponential phase (OD = 0.57−0.59), were diluted back to OD = 0.02, and grew again to saturation (OD = 2.8−3.1). Thus the cells grew in minimal media for a total of about 12 generations. To compare the abundance of each strain at the end of each experiment to its abundance at the beginning, we used DNA barcode sequencing (42) with Illumina. Specifically, we extracted genomic DNA and performed PCR using the 98°C protocol (17).

We sequenced these three samples using Illumina HiSeq. For each sample, we obtained 23-26 million reads with barcodes that matched the pool. We also sequenced these three samples on a MiSeq instrument, along with two additional samples that were collected from the two replicate cultures before the first transfer (at about 5 generations). The MiSeq run had 1.7-3.0 million reads per sample.

We computed gene fitness values as described previously (17). Briefly, the fitness of a strain is the normalized log_2_ ratio of the number of reads, and the fitness of a gene is the weighted average of the fitness values for strains with insertions in the central 10-90% of the gene. In each HiSeq sample, the median gene had around 2,000 reads for relevant strains. The two replicate cultures yielded similar gene fitness values (r = 0.995; Figure 3A). Fitness at 12 generations (from HiSeq) was strongly correlated with fitness at 5 generations (r = 0.936, Figure 3B). Gene fitness at 12 generations was also similar (r = 0.91) to the results of an independent experiment on a different day with the same pool of mutants, a higher concentration of D-glucose (3.96 g/L), a smaller volume (10 mL), and fewer generations of growth (about 6.9; experiment set2IT096 of (17)).

**Figure 3.**
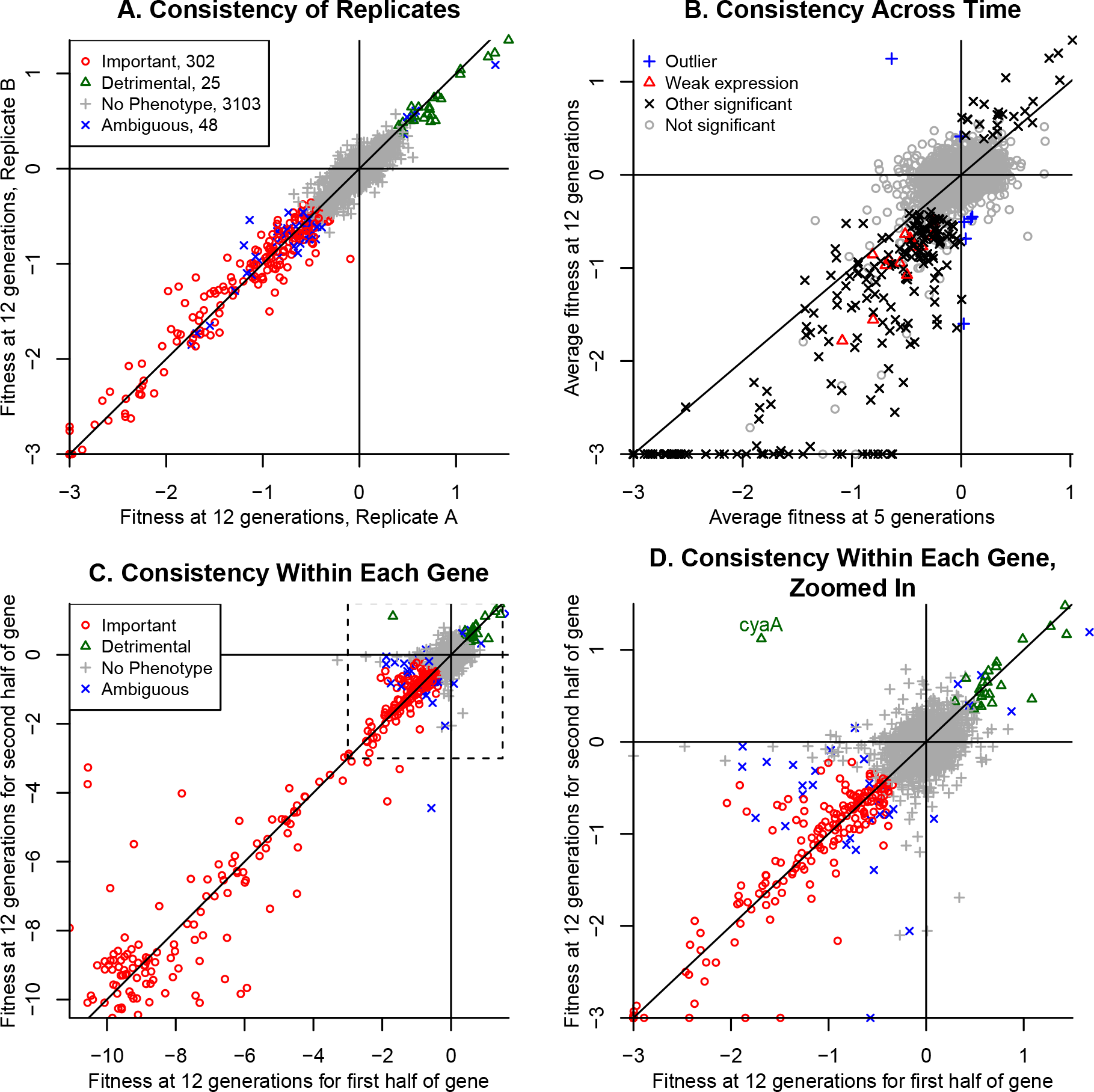
**Consistency of gene fitness values in minimal glucose medium.** (A) Consistency between replicates at 12 generations. Fitness values less than −3 are shown at −3, and 123 genes have fitness under −3 in both replicates. (B) Consistency across time. Genes with significant phenotypes (of either sign) are subdivided into those with weak expression (under 2 ppm of monomers) or above. Fitness values less than −3 are shown at −3. (C) Consistency within each gene. Fitness values at 12 generations were computed separately for the first and second half of each gene that had sufficient coverage. (D) shows the same data as (C), but only for fitness values above −3. In all panels, lines show *x* = 0, *y* = 0, and *x* = *y*.

Gene fitness values at 12 generations were usually more extreme than those at 5 generations, which indicates that the abundance of the mutants continued to change in the same direction from 5-12 generations as they had during 0-5 generations (Figure 3B). This shows that most of these genes were important during exponential growth, which is when the ribosomal profiling data was collected, and not just for the transition to growth in this medium. All 13 genes that were significantly important for fitness (as described below) despite weak expression of under 2 ppm of monomers were also below the line (Figure 3B). (None of the genes that were detrimental to fitness were so weakly expressed.)

As another way to check that the fitness data represents the impact of disabling each protein, we estimated the fitness at 15 generations separately for the first and second half of each gene, and then averaged the two independent replicate experiments (Figure 3C). We considered the 3,189 non-essential proteins that were included in the comparison to the ribosomal profiling data and have sufficient coverage of both halves of the gene. If the change in abundance of the strains is due to disabling the protein, then insertions at different locations in the gene should have similar fitness values, while if the change in strain abundance is due to some other factor such as secondary mutations, then the insertions at different locations would have uncorrelated fitness values. The fitness values from the two halves were strongly correlated (r = 0.96), and just one gene had a significant phenotype but had inconsistent signs for the two halves (the adenylate cyclase *cyaA*, see Figure 3D).

A caveat that our analysis did not take into account is polar effects, in which a transposon insertion in one gene leads to reduced expression of downstream genes via rho-dependent termination. Although we do not believe polar effects are common (see (17) for a systematic analysis), they might lead us to underestimate the proportion of “useless” protein.

### Identifying significant phenotypes in minimal glucose medium

For each replicate and for each gene, we computed a *t*-like test statistic that takes into account the variability of the fitness values for the strains for each gene (17). To identify genes with statistically significant changes, we wanted to combine the two *t* values, but they are not independent as they both used the same data for the initial sample. So, instead of using the usual way of combining *t* values of (*t_A_* + *t_B_*)/√2, we used *t_comb_* = (*t_A_* + *t_B_*)/√3. The increased denominator makes up for the partial nonindependence. To see why 3 instead of 2 is correct, consider the expected variance of the sum of the two fitness values, and remember that fitness is the difference of log abundances. With non-independent start samples, the variance is Variance(*A* — *C* + *B* — *C*) = Variance(*A*)+Variance(*B*)+4· Variance(C), while with independent start samples, it is Variance(*A* — *C* + *B* — *D*) = Variance(*A*)+Variance(*B*)+Variance(C)+Variance(D). (Remember that Variance(*A* + *B*) = Variance(*A*) + Variance(*B*) if *A* and *B* are independent.) If each component has about the same variance, then the non-independence increases the variance by a factor of 6/4 = 3/2.

Genes were considered to significantly affect fitness if |*t_Comb_*| > 4. If *t* follows the standard normal distribution then we expect 0.2 false positives. As a control, we compared the two 12-generation samples to each other and identified just 1 gene with |*t*| > 4.

Seven genes had strains with statistically significant changes at 12 generations but had a different sign after 5 generations (these can be seen as outliers in Figure 3B). Another 39 genes had mutants that showed non-significant changes in abundance of 3% per generation or more. These 48 genes were classified as ambiguous.

Most genes whose deletion strains have strong growth defects when grown individually in glucose medium also have a significant fitness defect in our assay. Baba and colleagues (18) grew each mutant from the Keio deletion collection in a minimal MOPS glucose media and measured the optical density at 24 and 48 hours. (Although their media was similar to ours, it contained 2 mM instead of 1.32 mM phosphate.) We focused on the measurement at 24 hours as it should be more sensitive to a reduction in the growth rate, and we considered a 2-fold reduction in OD relative to the median as a strong defect. Of the genes that were included in the comparison to the ribosomal profiling data, 140 had such a strong reduction in OD, and in the fitness data, we classified 116 (83%) of these as significantly important for fitness, 13 as ambiguous, 1 as detrimental (*rseA*), and 10 as not important for fitness. *RseA* encodes an anti-sigma factor of σ^E^ and is not important for fitness in our other defined-media experiments either (17); the cause of the discrepancy is not clear. The other 10 discordant genes included several genes involved in molybdenum cofactor synthesis (*moeA, moeB, moaC*) or selenocysteine synthesis (*selD*), but these processes are not expected to be important for growth under aerobic conditions. The discrepancy probably indicates a low concentration of oxygen in the experiment of Baba and colleagues, which was conducted in 96-well microplates without shaking. The other discordant genes were *ubiC*, which is involved in ubiquinone synthesis but is not required for growth on glucose (43); the DNA repair enzyme *uvrD*; *exoX*, which encodes a genetically redundant DNA repair enzyme (44), so it is not clear why it would be important for fitness in these growth conditions; *metL*, which is expected to be redundant with *thrA* in these conditions; and the poorly characterized genes *hflD* and *ycaL*.

### Proteomics confirms the high expression of “unnecessary” proteins

To check that the high expression of genes with no measurable benefit is genuine, we considered proteomic estimates of protein abundance (15). Unfortunately, the proteomic data with the highest coverage is from cells that were grown in a different media formulation than we used (M9 media with trace elements added instead of MOPS media, and with a higher concentration of glucose). Nevertheless, among the 4,062 proteins that were included in our analysis, 1,909 were quantified by proteomics, and for these proteins, the proteomics data was quite correlated with the ribosomal profiling data. For example, the Spearman rank correlation of the weight fraction, according to the two different methods, was 0.84, and the Pearson linear correlation of the monomer fractions was 0.82. The proportion of expression (by weight) that was due to proteins with no measurable phenotype was 23% according to proteomics, as compared to 29% for these same proteins according to ribosomal profiling. (29% is slightly less than 31%, which was reported above, because of the aggregate expression of the proteins that were not quantified by proteomics.)

We also checked if the 287 highly-expressed “unnecessary” proteins (with a weight fraction of above 200 ppm) were also highly expressed according to the proteomics data. 270 of the 287 proteins (94%) were quantified by proteomics, and of these, 219 (81%) had a weight fraction of above 100 ppm in the proteomics data. Overall, the proteomics data confirmed the high expression of most of these proteins.

### Experimental differences between the fitness data and the ribosomal profiling data

The ribosomal profiling data was from strain MG1655, while our transposon mutants were made from strain BW25113. The genomes of these strains were compared by (45). BW25133 lacks *araBAD*, *rhaDAB*, or *valX*, has a truncated and modified *lacZ*, has a frameshift in *hsdR*, and has a premature stop codon in *yjjP*. BW25133 also lacks 110 nt in an intergenic region. MG1655 (but not BW25113) has mobile element insertions in *crl*, in *mhpC*, and in two intergenic regions, and has frameshifts in *glpR* and in *gatC*. The strains also differ at a 3 nt stretch in *rrlD* and 13 other singlenucleotide substitutions.

We do not expect these differences to lead to global changes in gene expression or in growth. Indeed, the proteomics data (15), which gave similar results as the ribosomal profiling data, was obtained using BW25113. Incidentally, both strains have a frameshift mutation in *rph* that reduces the expression of the downstream gene *pyrE*, which is required for pyrimidine biosynthesis; this mutation reduces the growth rate in minimal media by around 10% (46).

The media formulations for the fitness and ribosomal profiling experiments were identical, and both experiments used a culture volume of 200 mL at 37°C and shaking at 180 rpm. However, the ribosomal profiling experiments used 2.8 L flasks, while for mutant fitness experiments we used 2.0 L flasks, so the concentration of oxygen might not have been identical. The proteomics data used a different media and a smaller volume: it was collected using M9 media with 5 g/L glucose and added trace minerals, and with a 50 ml culture in a 500 ml flask shaking at 300 rpm (15).

### A fitness compendium for Escherichia coli K-12

This compendium includes previously-described fitness experiments with various carbon sources and M9 minimal media (17). Nitrogen source experiments were conducted similarly, with D-glucose as the carbon source. Stress experiments were conducted in LB, at 28^o^C instead of 37°C, and in a 48-well microplate. Inhibitors were added at a concentration that would reduce the growth rate by about 2-fold. All of these experiments were inoculated at OD = 0.02 and grown aerobically until saturation. Only experiments that met standards for internal and biological consistency (17) were retained for analysis.

Fitness values and *t* scores were obtained as described above. *t* values from replicate experiments were combined as described above if they shared a control. If there were 3 or 4 replicates with shared controls, then the variance was reduced by 2 or 2.5 instead of by 1.5 fold. If the replicates were fully independent, then the t values were combined in the traditional way (*t_Comb_* = −*t*/√*n* where *n* is the number of replicates). For stress experiments, only independent samples grown at the same concentration were considered to be replicates.

Across the compendium, genes were considered to have a significant phenotype if, in any condition, average fitness was under -0.5 and *t_Comb_* < −4. There were 1,107 such genes. In 13 control comparisons between independent samples from the same culture, this threshold was never reached. (Given 13 · 3, 789 values from the standard normal distribution, the expected number of values under −4 would be 1.6.) Based on the standard normal distribution, we would expect 13 false positives in this data set, or a false discovery rate of about 1%.

For the analysis of individual experiments, a phenotype was considered significant if fitness was under −1 and *t* < —4.

## Software

The *E. coli* fitness experiments were analyzed using FEBA statistics version 1.0.1 (https://bitbucket.org/berkeleylab/feba). Statistical analyses were conducted in R 2.15.0.

## Data Availability

Tables of counts per barcode, fitness values, and *t* values are available at http://genomics.lbl.gov/strongselection/, as are supplementary tables 1 and 2. Also, the *E. coli* fitness compendium can be browsed at http://fit.genomics.lbl.gov/.

## Appendix: Comments on the functions of highly-expressed and “unnecessary” proteins

70 of the 287 proteins are involved in key processes that we expected would be important for growth in minimal glucose media. For 60 of them, we can explain the lack of phenotype due to redundancy. The redundancy may be due to isozymes, but sometimes the redundancy is indirect. For example, SpeD is highly expressed and is required for the synthesis of spermidine, which is one of the major polyamines in *E. coli*. Previous studies found that a strain of *E. coli* that lacks spermidine has a subtle growth defect (around 15%) while a strain that lacks both spermidine and another polyamine, putrescine, has a severe growth defect (around 70%) (47; 48). This indicates that putrescine and spermidine synthesis are partly redundant, and under our conditions, spermidine synthesis may be fully redundant. Alternatively, the effect of the loss of spermidine might be small: SpeE is also required for spermidine synthesis, and we found that knockouts of *speE* had a subtle growth defect (3% per generation) that is near the limit of sensitivity of our fitness assay.

Why doesn’t mutating any of the other 10 highly-expressed proteins that are involved in key processes lead to reduced growth? BamB, BamC, SecB, and YajC are non-essential components of essential protein complexes. LepA, MiaB, and RimO are accessory proteins for translation or for the modification of tRNA or rRNA, and might have more subtle advantages. ZapB is required for Z ring placement and mutants have altered size but still grow at about the same rate as wild-type cells (49). TatA is the twin arginine translocase and apparently none of the proteins that it exports are important for growth in our conditions. And MlaC is involved in ensuring the asymmetry of lipids in the outer membrane; this is important for stabilizing the outer membrane but is not important for fitness in standard growth conditions (50).

## Supplementary Tables

Supplementary Table S1: Comparison of fitness data and ribosomal profiling data.

Supplementary Table S2: Manual classification of highly expressed genes with no phenotype in minimal glucose.

**Supplementary Figures**

**Figure S1.**
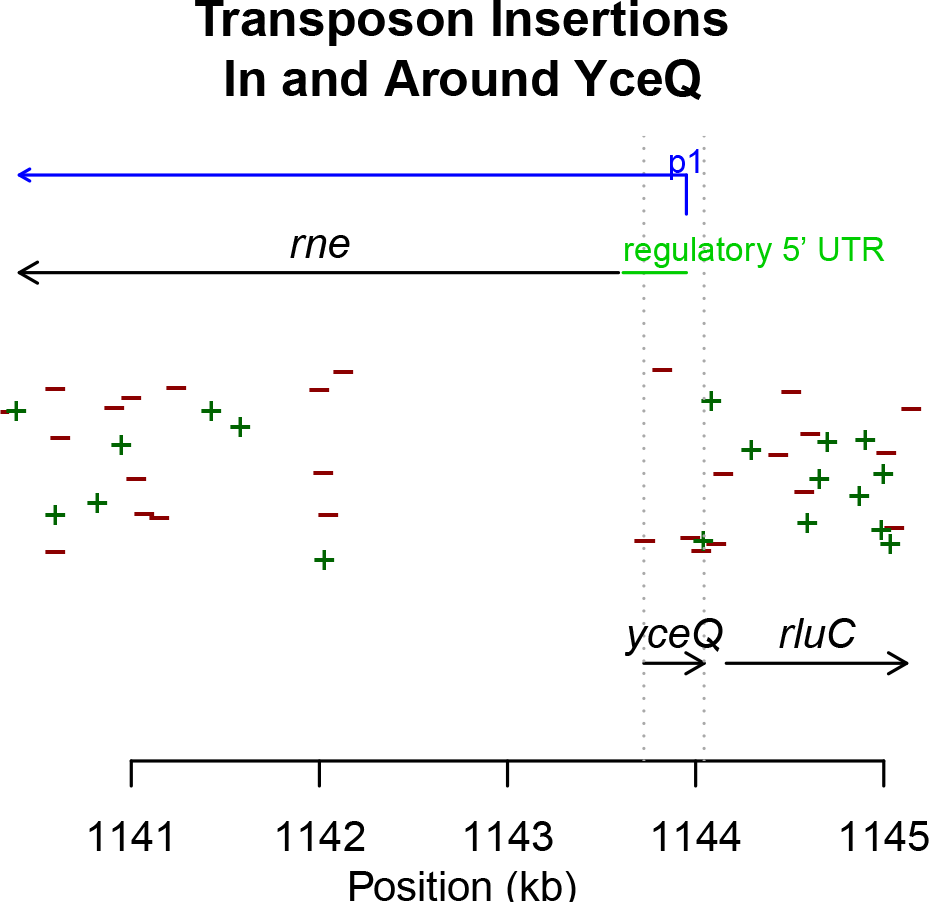
**Locations of transposon insertions in and around yceQ.** Only locations that are supported by at least two TnSeq reads (Wetmore et al 2015) are shown. A “+” symbol indicates that the promoter and the antibiotic resistance gene within the transposon are on the + strand. We also highlight the location of the primary promoter (p1) of *rne* (Ow et al, Molecular microbiology 43:159-171), which is an essential gene, and the conserved and structured regulatory leader in the 5’ untranslated region of the rne mRNA (Diwa et al, Genes & development 14:1249-60). The vertical lines show the extent of *yceQ*. Note that *yceQ* overlaps both the primary *rne* promoter and the regulatory leader, which may explain why deleting the entire region is not possible (Baba et al 2006). Consistent with this, we identified transposon insertions within *yceQ* but only in the “-” orientation, so that the promoter within the transposon could drive expression of *rne*. Incidentally, insertions within the 3’ part of *rne* are viable because the C-terminal part of RNase E is not required for its catalytic activity or for its essential role in processing ribosomal RNA (Kido et al, J. Bacteriology 187:3917-25; Lopez et al, Molecular microbiology 33:188-99).

## Acknowledgements

We thank Mark Callaghan for technical assistance with the *E. coli* fitness compendium.

